# Role of SCN5A coding and non-coding sequences in Brugada syndrome onset: What’s behind the scenes?

**DOI:** 10.1101/218099

**Authors:** Houria Daimi, Amel Haj Khelil, Ali Neji, Khaldoun Ben Hamda, Sabri Maaoui, Amelia Aranega, Jemni BE Chibani, Diego Franco

## Abstract

Brugada syndrome (BrS) is a rare inherited cardiac arrhythmia associated with a high risk of sudden cardiac death (SCD) due to ventricular fibrillation (VF). BrS is characterized by coved-type ST-segment elevation in the right precordial leads (V1-V3) in the absence of structural heart disease. This pattern is spontaneous, or is unmasked by intravenous administration of Class I antiarrhythmic drugs. The SCN5A-encoded α-subunit of the NaV1.5 cardiac sodium channel has been linked to BrS, and mutations in SCN5A are identified in 15–30% of BrS cases. Genetic testing of BrS patients generally involves sequencing of protein-coding portions and flanking intronic regions of SCN5A, according to recent international guidelines. This excludes the regulatory untranslated regions (5’UTR and 3’UTR) from the routine genetic testing of BrS patients. We here screened the coding sequence, the flanking intronic regions as well as the 5’ and 3’UTR regions of SCN5A gene and further five candidate genes (GPD1L, SCN1B, KCNE3, SCN4B, and MOG1) in a Tunisian family diagnosed with Brugada syndrome.

A new Q1000K mutation was identified on the SCN5A gene along with two common polymorphisms (H558R and D1819). Furthermore, multiple genetic variants were identified on the SCN5A 3’UTR, one of which is predicted to create additional microRNA (miRNAs) binding site for miR-1270. Additionally, we identified the hsa-miR-219a rs107822. No relevant coding sequence variant was identified in the remaining studied candidate genes. Although Q1000K is localized in the conserved binding site of MOG1 which predicts a functional consequence, this new mutation along with the additional variants were differentially distributed among the family members without any clear genotype-phenotype concordance. This gives extra evidences about the complexity of the disease and suggests that the occurrence and prognosis of BrS is most likely controlled by a combination of multiple genetic factors and exposures, rather than a single polymorphism/mutation. Most SCN5A polymorphisms were localized in non-coding regions hypothesizing an impact on the miRNA-target complementarities. In this regard, over-expression of miR-1270 led to a significant decrease of luciferase activity suggesting a direct role regulating SCN5A. Therefore, genetic variants that disrupt its binding affinity to SCN5A 3’UTR and/or its expression might cause loss of normal repression control and be associated to BrS.

## INTRODUCTION

Brugada syndrome is diagnosed when a type-1 ECG pattern is observed in the precordial leads (V1-V3), in the presence or absence of a sodium channel blocker agent, and in conjunction with one of the following features: documented ventricular fibrillation (VF), polymorphic ventricular tachycardia (VT), a family history of sudden death (SD) at an age younger than 45 years, the presence of coved-type ECG in family members, inducibility of ventricular arrhythmias with programmed electrical stimulation, syncope, or nocturnal agonal respiration [1].

Brugada syndrome is genetically determined, with an autosomic dominant pattern of transmission and predominantly affected males [2]. Approximately 60% of patients with aborted sudden death with the typical BrS electrocardiogram have a family history of sudden death, or have family members with the same electrocardiographic abnormalities, supporting a familiar inheritance of the disease. However, sporadic cases of BrS have also been reported [3, 4]. Genetic determinants of the BrS have emerged over the last decade [5]. Mutations in SCN5A, the gene encoding the cardiac sodium channel Nav1.5, account for 15-30% of clinically affected patients and around 400 mutations in the SCN5A gene were found in probands with BrS. Many of these mutations were functionally studied and showed a spectrum of biophysical abnormalities all leading to Nav1.5 (SCN5A) loss of function [6]. However, it was found that some large BrS-affected families contained SCN5A-positive and SCN5A-negative family members which questions the direct causative role of SCN5A gene in BrS [7].

Recently, seventeen susceptibility genes were identified for BrS. However, mutations in these genes account for only 35–40% of familial BrS [8]. Therefore, to date SCN5A remains the most frequently affected candidate gene which is fully associated with isolated BrS, yet despite so, the genetic bases of approximately 80% of the diagnosed BrS patients remains elusive.

More recent studies suggested that miRNAs play important roles in cardiovascular diseases and particularly cardiac arrhythmias [9]. miRNAs are short noncoding RNA molecules that regulate gene expression by binding with the 3’ UTR of their target genes, leading to the degradation or in some rare cases to the stabilization of their target’s mRNA [10, 11]. miRNAs are emerging as pivotal regulators of gene expression and protein translation involving processes such as cardiac remodeling, growth and arrhythmias [12, 13, 14, 15]. Thus, the loss or gain of function of a specific miRNA represents a vital event in heart diseases and might have a strong impact on the manifestation of inherited diseases such as BrS.

In the current study, we report a Tunisian family with BrS carrying a new SCN5A Q1000K mutation combined with two common SCN5A polymorphisms H558R and D1819D. Our findings revealed a heterogeneous distribution of genetic variants along the SCN5A gene sequence with no genotype-phenotype correlation, arguing that additive rather than a single defect underlies BrS phenotype. The absence of relevant genetic alterations in the most reported BrS candidate genes gives more evidences that BrS in this family is SCN5A associated. Interestingly, more than 70% of the identified SCN5A genetic variants are localized in the 3’UTR, supporting an important role of this region in the BrS phenotype.

## MATERIALS AND METHODS

### 1 Clinical studies/ study subjects

The current study relates to a Tunisian family followed up for more than five years at the Cardiology Unit in Monastir Hospital (Tunisia) after a confirmed Brugada syndrome first diagnosed in its proband **(II-1)**. The clinical follow-up included a review of personal and familial history, complete physical examination, 12-lead ECG, 24-hour Holter ECG monitoring, and echocardiography. The flecainide challenge test was performed for four family members **(II-3, III-1, III-2 and III-3)**. The proband’s mother and wife declined the clinical test however, a blood sample was obtained from all family members including both women for the genetic testing. An informed consent was obtained from all participants.

In the control group, individuals with cardiac diseases or any familial history of heart disease were excluded. As control cohort, 100 blood samples of healthy Tunisian individuals were obtained from the Banque de sang-Monastir-Tunisia. Age ranges from 35 to 75 years ±15. The study protocol was approved by the Human Ethics Committee of both Monastir University School of Medicine–Tunisia and University of Jaén-Spain. Blood samples were processed as detailed in **Supplementary data**.

### 2 Genetic screening

Genetic screening for mutations in SCN5A gene was performed by PCR-based Sanger sequencing (Sistemas Genómicos, Valencia, Spain). PCR amplification was performed using originally designed oligonucleotide primers encompassing all 28 exons, ~150-bp adjacent intronic areas, 3’UTR and 5’UTR. Chromatogram files were processed using *BioEdit* software v7.2.6.1 (Caredata.com, Inc) and sequences were aligned against consensus sequences using Blast (https://blast.ncbi.nlm.nih.gov). Genetic variants were validated after three rounds of sequencing. The same protocol was applied to screen for mutations the following genes: GPD1L (NM_015141.3), SCN1B (NM_199037), KCNE3 (NM_005472.4), SCN4B (NM_174934.3) and MOG1 (NM_001177802.1).

All mutations on SCN5A gene were denoted using known and accepted nomenclature based upon the full-length of the splice variant with 2,016 amino acids (aa) (PubMed Accession No. NM 198056). The location of each domain (D) of NaV1.5, was predicted using Swissprot (http://ca.expasy.org/uniprot/)

### 3 Cell culture and transfection assays

HL-1 cells were cultured as described by Claycomb et al. [16] and incubated at 37°C in a humidified atmosphere containing 5% CO_2_. Cells were plated in six well cell culture dishes and transfected with pre-miRNA-1270 as well as with negative controls, such as FAM-labeled control pre-miR, scrambled pre-miR or non-transfected cells following supplier’s guidelines (Ambion). FAM-labeled premiR negative control transfected cells allowed evaluating the transfection efficiency, which it was in all cases greater than 50%, as revealed by fluorescence microscopy recording. Cells were collected 6 hours after transfection.

### 4 RNA isolation and qRT-PCR

Negative control and transfected cells were collected and a Trizol-base RNA isolation was performed according to the standard protocol as detailed in Supplementary data. *Scn5a*, *Mef2c* and *Nkx2.5* expression levels were qRT-PCR quantified on every transfection condition. Data analysis was performed as described by Livak & Schmittgen [17].

### 5 Luciferase Reporter Constructs and assays

pMIR-Report luciferase SCN5A-3’UTR reporter vector was generated as detailed in Supplementary data. LC5 human lung fibroblasts (Vircell, Granada-Spain) were plated transfected with SCN5A 3’UTR-luciferase or an empty vector, respectively. After 24 hours of transfection, cells were incubated with 50nM of the corresponding pre-microRNAs (Ambion) and cultured for another 24 hours. Cells were collected and luciferase activity was measured as detailed in Supplementary data. Experiments were repeated at least three times for further validation.

### 6 Statistical analyses

Statistical analyses were performed using a t-student test. A p<0.05 was considered statistically significant.

## RESULTS

### 1 Clinical analysis

The index case (**Subject II-1, Figure 1A**) is a 51-year-old man, presented to the Cardiology service in Mahdia Hospital (Central Tunisia) after three successive episodes of palpitation associated with dizziness and lipothymia with a normal resting ECG recording. Transferred thereafter to Marsa Hospital-Tunisia, the patient received a transthoracic echocardiography, an exercise ECG testing and coronarography all without any abnormalities. The 51-year-old man remained asymptomatic for about two years since this incident. However, during a flight; he presented palpitations again with dizziness. The proband was admitted in Frankfurt Hospital (Germany) where he was submitted to electrocardiographic and coronarographic essays, but again all tests resulted normal.

**Figure 1:**
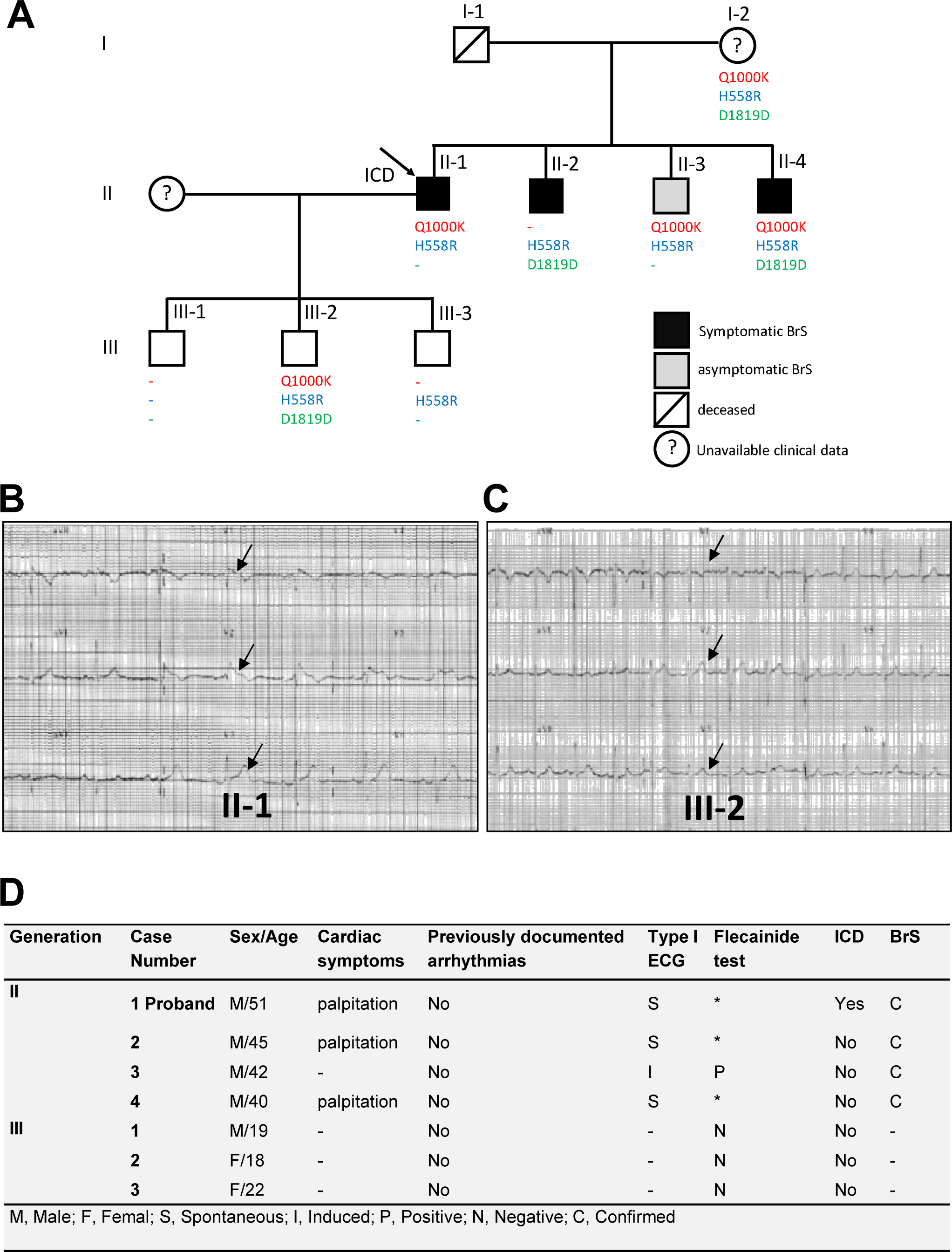
Family kindred and mutations/polymorphisms distribution. **Panel A:** Family kindred. Q1000K, H558R and D1819D distribution is displayed, illustrating a lack of genotype-phenotype correlation. Black cells refer to symptomatic BrS patients presenting with spontaneous BrS type 1 ECG, Gray cells refer to asymptomatic patients presenting with induced BrS type 1 ECG and white cells refer to patient presenting no BrS symptoms nor type 1 BrS ECG. **Panels B and C:** 12 leads ECG recorded from II-1 and III-2 patients respectively. Note the ST segment elevation in the precordial leads V1, V2 and V3 in II-1 patient ECG. Both II-1 and III-2 patients are carriers of the Q1000K mutation. However, N-III-2 patient presents no symptoms for BrS nor Type 1 ECG which could be due to its young age and/or the incomplete penetrance of the disease. **Panel D:** Clinical profiles of subjects belonging to Family N. Note that there is a heterogeneous clinical profile within the symptomatic patients. M: male; F: Female; S: Spontaneous; *: Not applicable; N: Negative; P: Positive; C: Confirmed; I: Induced.

Few months later, the proband was admitted again in Monastir Hospital–Tunisia after palpitation episodes associated with dizziness and perfuse sweating without loss of conscience. The patient presented elevated arterial blood pressure with ventricular tachycardia at 220 beat/min. He was then applied to two external electrical shocks returning thereafter to the sinusal rhythm at 70 beat/min. 1mg/kg of Xylocaine (class 1b antiarrhythmic) was administered followed by continuous infusion at 1,5mg/kg/h. This unmasked an ST segment elevation of 3mm in V2, V3 and V4 and an incomplete right bundle branch block confirming a BrS diagnosis (**Figure 1B)**. An implantable cardioverter defibrillator (ICD) was implanted to the BrS patient few weeks later.

The clinical investigation of the whole family revealed a heterogeneous diagnosis classifying the family patients into three principal categories i.e. symptomatic, asymptomatic and no BrS patients. Symptomatic patients refer to those presenting aborted sudden cardiac death or specific symptoms (syncope, palpitations and/or dizziness) and displaying type 1 (coved-type ST segment elevation ≥ 0.2 mV followed by a negative T wave) BrS ECG (either spontaneously or after sodium channel blockade) whereas, asymptomatic patients refer to those presenting no specific symptoms but to whom a Brugada type ECG was unmasked after a flecainide challenge test. No BrS subjects refer to those presenting neither specific symptoms nor type 1 BrS ECG even after flecainide test (**Figure 1C and D)**. No spontaneous Brugada type ECG was recorded in the asymptomatic patients belonging to this family neither a cardiac conduction defects was detected.

### 2 Genetic analysis of SCN5A and candidate genes

Mutation screening of SCN5A gene identified two widely studied common polymorphisms: H558R and D1819D **(Figure 2B, C)**. H558R (Exon-12, c.1673 A>G, rs1805124) polymorphism was identified in all family members except III-1 patient, while D1819D (Exon-28, c.5457 C>T, rs1805126) was detected in four members of the family. The prevalence of H558R and D1819D in the general population is 0,224 and 0,553 respectively. In addition to both polymorphisms, a new missense variant c. 3192 C>A was revealed in Exon 17 leading to the p.Q1000K substitution (**Figure 2A**). This variant was not found in a control group of 100 ethnically matched healthy controls. Furthermore, this amino acid change is not reported in different publically available SNP databases (1000 genome, NHLBI Exome variant server and NCBI) in which thousands of control samples have been sequenced, demonstrating thus that such missense variant corresponds to a new mutation. Q1000k missense variant was revealed in the proband (II-1) in addition to four members of his family (I-2, II-3, II-4 and III-2) **(Figure 2C)**. Surprisingly, we did not find this variant in the proband’s brother (II-2) who is a symptomatic BrS patient while this variant was revealed in the proband’s son who is negative for the disease (III-2).

**Figure 2:**
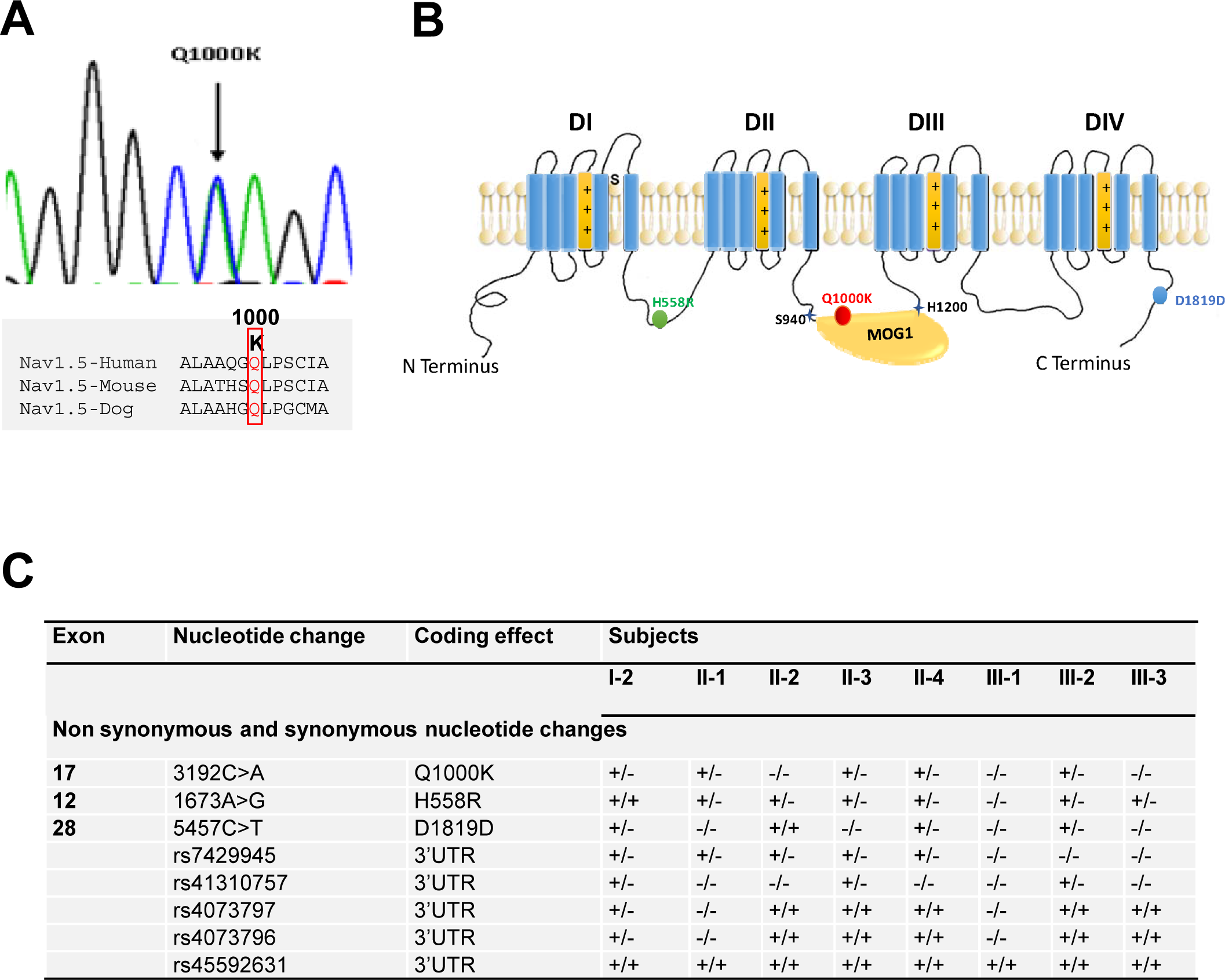
SCN5A nucleotide changes among the family members. **Panel A** A new missense variant c. 3192 C>A was revealed in Exon 17 leading to the p.Q1000K substitution. Q1000 is conserved among species which may predict a crucial functional impact of Q1000K mutation **Panel B** Localization of the three identified amino acid changes among the whole Nav1.5 protein structure. Note that Q1000K is localized in the conserved binding site of MOG1 which predicts a functional consequence. **Panel C** Summary of the SCN5A mutations and polymorphisms distribution among the family members.

Genetic analysis of SCN5A promoter region and 5’ UTR failed to reveal any genetic variation. However, within the 3’UTR, five distinct previously reported polymorphisms were identified: rs7429945 (c.*123A>G), rs41310757 (c.*753C>T), rs4073797 (c.*962T>A), rs4073796 (c.*963C>T) and rs45592631 (**Figure 2C**).

Bioinformatics analysis demonstrated that 3’UTR genetic variants can affect distinct microRNAs binding sites. In this context, rs4073797 and rs4073796 polymorphisms create a new miR-1270 binding site (position 962-969 of SCN5A 3’UTR) (**Figure 3A**), in addition to other three poorly conserved miR-1270 binding sites already predicted within the SCN5A 3’UTR (Position 713-719, position 1824-1830 and position 1852-1858 of SCN5A 3' UTR). It is plausible that by altering the genetic sequence of SCN5A 3’UTR, miR-1270 might increase its binding affinity to the SCN5A 3’UTR resulting therefore in impairing SCN5A transcript bioavailability. Thus, using a pMIR-Report luciferase SCN5A 3’UTR construct we demonstrated that miR-1270 over expression significantly decrease luciferase activity as compared to pMIR-Report luciferase controls (**Figure 3C**). In order to demonstrate if similar effects were observed in a cellular context, HL-1 atrial cardiomyocytes were transfected with miR-1270. miR-1270 overexpression displays a significant reduction on Scn5a expression, in line with the luciferase assays (**Figure 3B**). Specific targeting to Scn5a is demonstrated since miR-1270 does not influence Nkx2.5 and/or Mef2c expression in HL-1 cells (**Figure 3B**) confirming that genetic variants creating additional binding sites for miR-1270 (rs4073797 and rs4073796) can imbalance Scn5a expression.

**Figure 3:**
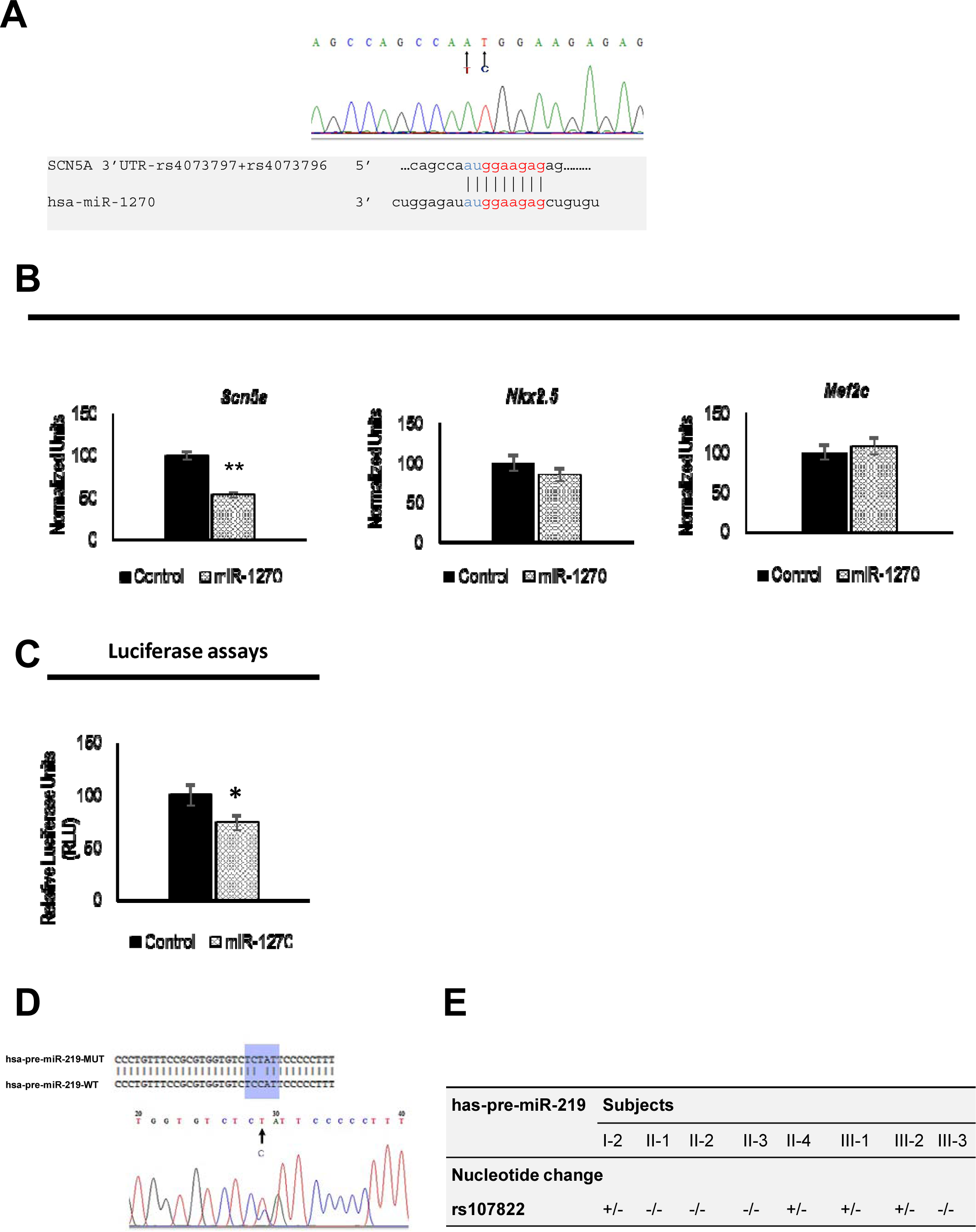
SCN5A 3’UTR nucleotide changes and impact on micrRNAs binding sites. **Panel A** Sequencing results of exon 28 reveals two already reported successive polymorphisms T83948A and C83949T. Together, these polymorphisms create a forth miR-1270 binding site within SCN5A 3’UTR as detailed. **Panel B** Over-expression of miR-1270 in HL-1 cardiomyocytes demonstrated a significant decrease of Scn5a mRNA quantity (* p<0.05, **p<0.01) 6 hours post-transfection. qRT-PCR of two control genes Nkx2.5 and Mef2c in HL1 cells over-expressing miR-1270 displayed no difference of expression in transfected cells compared to control condition. Black bars refer to control condition (non-transfected cells) and patterned bars to the experimental condition (transfected cells). **Panel C** qRT-PCR results are also confirmed by luciferase assays as miR-1270 gain of function decrease the luciferase activity in cultured human fibroblasts. **Panel D** Sequencing of miR-219a precursor and flanking regions identified the heterozygous alleles of rs107822 SNP which is a substitution C>T localized 36 bp upstream the miR-219a precursor sequence. **Panel E** Distribution of rs107822 nucleotide change among the family members.

In order to further explore the genetic background of BrS in this family specially in the Q1000K-negative patients, additional genetic screenings were performed for GPD1L, SCN1B, KCNE3, SCN4B and MOG1. Only previously described GPD1L (c.*1029A>G, rs6802739) and SCN4A (c.*1820G>A (rs45539741) and c.*3054T>C (rs3741315)) 3’UTR polymorphisms were identified (**Supplementary Figure 1**), however, without any colocalization with any regulatory feature mainly miRNAs binding sites.

### 3 Genetic screening of evolutionarily conserved microRNA genes targeting SCN5A

Recent studies have revealed that microRNAs (miRNAs) play an indispensable role in various facets of cardiac function through their alteration of target mRNAs expression [13, 14]. SCN5A is regulated either directly or indirectly by multiple microRNAs, i.e. miR-125a, miR-200a/b/c, miR-153a, miR-219-1, miR-98 and miR-106 as predicted by TargetScan (http://www.targetscan.org) and Pictar (www.pictar.mdc-berlin.de) algorithms and previously demonstrated by our group [11]. We therefore have screened the precursor sequences and ±300bp upstream and downstream flanking regions of all the microRNAs predicted to target SCN5A, in search of any genetic variants in these regions.

Genetic screening of miR-125a, miR-200a/b/c, miR-153a, miR-98 and miR-106 did not reveal any genetic variant. However, the rs107822C > T mapped 36bp upstream miR-219a precursor sequence was identified **(Figure 3D)**. This genetic variant was carried by four members (I-2, II-4, III-1 and III-2) (**Figure 3E**). Genetic screening of this variant in a control cohort of 100 individuals without cardiac defects demonstrated that rs107822C > T polymorphism was present in 11% of the cases (p<0.05). The physical localization of this SNP 36bp upstream the miR-219a rises the hypothesis that rs107822C > T might lead to impaired expression of miR-219a, therefore deregulating SCN5A/Nav1.5 expression. However, the lack of a complete co-segregation of C>T substitution with the Brugada syndrome besides its presence in the control cohort, means that C>T substitution could be rather considered as one of the molecular factors contributing to the genetic background of Brugada syndrome within this family rather than the causing effect of the disease.

## DISCUSSION

Our study relates to a Tunisian family with several members having BrS type 1 ECG either recorded spontaneously or unmasked after Flecainide challenge test. Available family members were submitted to genetic testing using a candidate gene approach screening the main potential genes recently described in association with the disease. In the present study, we identified one novel Q1000K (Exon 17, c. 3192 C>A) missense variant and two common polymorphisms (H558R and D1819D) in the SCN5A coding sequence. The Q1000K is absent in 100 healthy controls hypothesizing thus a new mutation. Q1000K mutation affects the intracellular segment connecting homologous domains II-III of Nav1.5. The amino acid properties analysis of Q/K demonstrates that Q1000K mutation leads to the change of an uncharged polar residue by a positive polar residue. Thus, such differences between both residues might lead to functional changes of the cardiac sodium channel. Furthermore, Q1000K mutation affects the intracellular segment connecting homologous domains II-III of Nav1.5 and is encompassed within the MOG1 binding domain which lays between the amino acids S994 and H1200 as described by Wu et al [18]. MOG1 is involved in the trafficking of Nav1.5 to the cell membrane by a direct protein-protein interaction [19]. This suggests a functional impact of Q1000K on the Nav1.5-MOG1 interaction. A plausible scenario suggests that Q1000K might reduce the affinity of MOG1 to Nav1.5 reducing thus the channel number at the cell surface which in turn leads to a decrease of I_Na_ currents [19]. It is noteworthy that mutations in the near neighboring amino acids of Q1000 such as A997T were described in association with a dramatic decrease of the cardiac sodium currents and were associated to Brugada syndrome which further reinforce our hypothesis [20, 21]. However, further functional assays are still to be performed in order to identify the real impact of Q1000K mutation. Interestingly, the Q1000K mutation was absent in the proband’s brother who is a symptomatic BrS patient while this variant was revealed in a healthy son of the proband which gives further evidences about the incomplete penetrance of SCN5A mutations in the Brugada syndrome context. Our findings are in agreement with previous studies demonstrating the difficulties in interpreting the results of a family-based genetic screening and underlining the phenotypic variability of *SCN5A* mutations [22].

In addition to Q1000K mutation, two SCN5A common polymorphisms have been identified: H558R (Exon-12, c.1673 A>G, rs1805124) and D1819D (Exon-28, c.5457 C>T, rs1805126). Single nucleotide polymorphisms have been shown to modify SCN5A mutations by either aggravation of the channelopathy or rescue of the pathophysiologic effect of the mutation [23]. H558R is one of the most common SCN5A-polymorphisms, with an allelic frequency of 20% to 30% within individuals of white ethnicity [24]. In the general population, the presence of SCN5A-H558R (without any SCN5A mutation) has been reported to be associated with increased heart rate, longer QTc intervals, and an increased susceptibility to atrial fibrillation in population studies [25]. Of significance, the minor R allele of this polymorphism, has been shown to alter SCN5A function by reducing depolarizing sodium current [26] and modulating the biological effects of concomitant SCN5A mutations [27,28] In the current study, H558R was revealed in a heterozygous allele in all the family members except the subject III-1. Only five patients have the H558R polymorphism combined with the new Q1000K mutation among them a healthy subject. Interestingly, the three Brugada syndrome negative subjects in this family are younger than 22 years, including the carriers of Q1000K/H5558R. While the subjects older than 40 years are all positive for the disease including patient II-2 who is negative for Q1000K. This rises two important facts shaping the Brugada syndrome physiopathology. The first fact is related to the impact of aging in the context of Brugada syndrome which could be partially explained by the proven decline of the cardiac sodium channel expression with age [29]. Although the role of aging on genetic variants in SCN5A remains unclear, it is reasonable to speculate that decreased sodium channel caused by aging will lead to a more serious phenotype for SCN5A mutation, which is somehow similar to what we observed in this family. The second fact is rather related to the complexity of the Brugada syndrome genetic background determination even in presence of SCN5A mutation. In this regard, we think that an additive rather than a single variant effect is what really shapes the BrS syndrome.

In addition, genetic variants such as synonymous polymorphisms that were belief as nonfunctional for decades might have hidden effects that could be finally unmasked with the recent advances in molecular biology. This is applicable to the D1819D polymorphism (rs1805126) which was identified in members of the study family. This SNP, located in the terminal coding region of the SCN5A gene, is a synonymous variant and has been dismissed as potentially causal. However, Spengler et al, [30] recently demonstrated that this polymorphism may modulate a nearby miR-24 site. Our *in silico* analysis demonstrated that miR-24 has two predicted poorly conserved binding sites in the SCN5A 3’UTR (Position 975-982 and position 1014-1021 of SCN5A 3' UTR). Thus, the alteration of miR-24 expression may alter the cardiac sodium channel expression. Moreover, this common genetic variant could be a genetic modifier of many SCN5A mutations known to cause Brugada syndrome [31] and it has been associated to a high risk of subsequent cardiac arrhythmias [32].

MicroRNAs (miRNAs) are small regulatory non-coding RNAs with important roles in a variety of physiological and pathological processes [33]. Among others they are instrumental in regulating numerous cellular processes associated with cardiac remodeling and disease, including arrhythmias [9]. We therefore explored if genetic variants in non-coding region of SCN5A and/or in microRNAs predicted to modulate SCN5A could be associated with BrS. Importantly, 5 out of 8 variants identified in this family are localized in the SCN5A 3’UTR, supporting a functional role of noncoding sequences on the genetic basis of BrS. We have demonstrated through bioinformatics analyses that the presence of rs4073797 and rs4073796 polymorphisms in the SCN5A 3’UTR creates a new binding site for miR-1270. Our functional assays in HL1 cardiomyocytes demonstrate that miR-1270 overexpression decreases Scn5a expression is in line with the presence of three predicted conserved binding sites for miR-1270 within the SCN5A 3’UTR (TargetScan www.targetscan.org and Pictar www.pictar.mdc-berlin.de). rs4073797 and rs4073796 polymorphisms create a fourth miR-1270 binding site and thus, we hypothesize that genetic variants creating a new binding site for miR-1270 in the 3’UTR further decrease Scn5a expression and thus contribute to the genetic bases of BrS.

As with any genomic sequence, miRNAs are prone to nucleotide variations that may have nonnegligible effects. The presence of a single nucleotide polymorphism (SNP) in the long miRNA primary (pri-miRNA) may affect its maturation process, its expression or the binding of the mature form to its target, which would then influence the expression of the target genes. We here have identified the rs107822 which is an allele T/C alternated polymorphism located in flanking sequence-36 bp of pre-miR-219a. Since our previous findings demonstrated that miR-219a over expression severely increases Scn5a expression [11], so we hypothesized that rs107822 within pri-miR-219a would affect its structure or expression, leading to an impaired miR-219a expression (most likely decreasing) which in turn will lead to a decreased SCN5A expression. Recently, Song et al. demonstrated that the miR-219a rs107822 GA and AA genotypes decreased the mature miR-219a expression compared with the miR-219a rs107822 GG genotypes in normal tissues which further supports our hypothesis [34] Although the functionality of this polymorphism has been studied in several pathological contexts including cancers and psychological disorders [35, 36], to our knowledge, it’s the first time that the rs107822 is identified in patients with cardiac diseases particularly BrS.

Importantly, the genetic variants with impact on microRNA binding affinity do not co-segregate with the BrS in affected members, supporting an additive contribution to the onset of BrS rather than a unique causative link, which further support our findings in the SCN5A coding sequence. Taken together, our results strengthen previous reports where genotype-phenotype relationship for BrS patients does not fully correlates with single genetic defect [37].

In summary, we were able to show a cooperative rather than a single causative variant as responsible for BrS syndrome. Furthermore, defects in non-coding regions of candidate genes and/or in their post-transcriptional modulators (microRNAs) should also be considered for genetic screening and functional studies in Brugada syndrome as well as probably in other arrhythmogenic settings.

## ACKNOWLEDGEMENTS

We would like to thank the Spanish National DNA Bank (BNADN, Salamanca) and the Tunisian National Blood bank (Banque Nationale de Sang) for their valuable support on DNA control samples (grant AL-09-0026). A special thank for the BrS patients participating in this study.

